# Sleep as a random walk - A superstatistical analysis of EEG data across sleep stages

**DOI:** 10.1101/2021.06.25.449874

**Authors:** C. Metzner, A. Schilling, M. Traxdorf, H. Schulze, P. Krauss

## Abstract

In clinical practice, human sleep is classified into stages, each associated with different levels of muscular activity and marked by characteristic patterns in the EEG signals. It is however unclear whether this subdivision into discrete stages with sharply defined boundaries is truly reflecting the dynamics of human sleep. To address this question, we consider one-channel EEG signals as heterogeneous random walks: stochastic processes controlled by hyper-parameters that are themselves time-dependent. We first demonstrate the heterogeneity of the random process by showing that each sleep stage has a characteristic distribution and temporal correlation function of the raw EEG signals. Next, we perform a superstatistical analysis by computing ‘hyper-parameters’, such as the standard deviation, kurtosis and skewness of the raw signal distributions, within subsequent 30-second epochs. It turns out that also the hyper-parameters have characteristic, sleep-stage-dependent distributions, which can be exploited for a simple Bayesian sleep stage detection. Moreover, we find that the hyper-parameters are not piece-wise constant, as the traditional hypnograms would suggest, but show rising or falling trends within and across sleep stages, pointing to an underlying continuous rather than subdivided process that controls human sleep.

## 1 Introduction

Many complex systems, such as the earth crust, the weather, biological organisms, or the stock market, show continuous fluctuations of their internal state variables, even in the absence of external perturbations. The underlying processes can often be quantified in the form of multivariate time series, and a mathematical analysis of the time series can be used to predict future states of the system, or simply to better understand its internal dynamics [1, 2, 3].

While in simple physical systems, state variables fluctuate around a fixed mean value and with a fixed variance (as in the case of local pressure variations in a gas at equilibrium), complex systems often have multiple dynamical attractors [4], that is, a set of qualitatively different modes of behaviour, between which the system will occasionally switch. Such transitions typically show up in the time series by a sudden (or gradual) change of the statistical properties of the fluctuating state variables.

A typical example of such mode switching behaviour is the sleep cycle in humans and other mammals, where the brain is passing through a sequence of seemingly distinct sleep stages [5]. In this case, multi-channel electroencephalographic (EEG) recordings offer a convenient way to quantify the ongoing changes in the brain over long periods, but also with high temporal resolution [6, 7]. The momentary amplitudes of an N-channel EEG recording represent a point in an N-dimensional state space, and the ongoing time-series of vectorial amplitudes defines a random walk within this high-dimensional space.

It is then a natural hypothesis that each sleep stage corresponds to a different cluster within EEG state space, and that the trajectory of the random walk moves to the corresponding cluster whenever a new sleep stage is entered. Indeed, we have confirmed this hypothesis in former work [8], where we applied a previously developed new method for analyzing and comparing spatiotemporal cortical activation patterns [9]. In this context, we have also analyzed the micro-structure of cortical activity during sleep and found that it reflects respiratory events and the state of daytime vigilance [10]. Moreover, we have developed a general method to quantify the separability of point clusters in high-dimensional state-spaces [11].

In principle, the existence of sleep-stage related clusters within EEG state space could be exploited for an automatic detection of these stages, based only on the momentary multi-channel amplitudes, or on short-time averages of those. However, modern methods of automatic sleep-stage detection are usually based on sliding time windows of a larger widths, so that the algorithm can also make use of temporal features in the EEG data that are characteristic for different sleep stages (such as sleep spindles or K-complexes) [12, 13]. In this case, it is not even necessary to record a large number of EEG-channels. Indeed, we have shown that reliable sleep-stage detection is even possible based on a single channel [14], thanks to the remarkable ability of machine learning systems to extract those features from the data that are most relevant for the classification task.

In this work, we continue our investigation of single-channel EEG data during human sleep. However, our present focus is not on the further improvement of automatic sleep stage detection, but on a more fundamental description of the statistical properties of EEG data, seen as a temporally heterogeneous random walk. In particular, we investigate how the random walk’s momentary statistical properties, also called ‘hyper-parameters’, are changing during and across sleep stages. Our approach is based on the novel method of superstatistical analysis [15, 16, 17], which we have originally developed to analyze the random migration patterns of individual cancer cells [18], revealing that their average migration speed, the directional persistence of the cell trajectories and other hyper-parameters are time-dependent, reflecting internal mode changes such as the cell cycle. In subsequent work, we have demonstrated that the method can also be used to extract and model gradual or abrupt hyper-parameter changes in other complex dynamical systems, such as the climate or the stock market [19].

In the present study, we apply a simplified version of superstatistical analysis to a set of full-night EEG recordings. Each 30-second epoch of these recordings has been visually scored by a sleep specialist, according to the AASM (American Academy of Sleep Medicine) rules, so that the data is categorized into 4 different sleep stages (REM, N1, N2, N3) and the wake state. For each sleep-stage-labeled epoch, we compute from the single-channel recordings certain statistical hyper-parameters, such as the standard deviation, the kurtosis, and the skewness of the EEG amplitude distributions. We show that also these hyper-parameters have characteristic, stage-dependent distributions which can be used for a simple Bayesian sleep stage detection. Moreover, we find that the hyper-parameters are not piece-wise constant, as the traditional hypnograms would suggest. Interestingly, they show rising or falling trends also within each of the sleep stages, pointing to an underlying continuous neural process that controls human sleep.

## 2 Methods

### Generation of data sets

This work is based on 68 independent data sets, each containing one full-night three-channel EEG recording (channels F4-M1, C4-M1, O2-M1) from a different human subject during sleep, recorded with a sampling rate of 256 Hz. For most of the following analysis, each of the three channels was treated as a different (sub-)data set and evaluated separately, except for the computation of the cross-correlation functions (see below). The participants of the study included 46 males and 22 females, with an age range between 21 and 80 years. Exclusion criteria were a positive history of misuse of sedatives, alcohol or addictive drugs, as well as untreated sleep disorders. The study was conducted in the Department of Otorhinolaryngology, Head Neck Surgery, of the Friedrich-Alexander University Erlangen-Nürnberg (FAU), following approval by the local Ethics Committee (323–16 Bc). Written informed consent was obtained from the participants before the cardiorespiratory polysomnography (PSG). After recording, the raw EEG data were analyzed by a sleep specialist accredited by the German Sleep Society (DGSM), who removed typical artifacts [20] from the data and visually identified the sleep stages in subsequent 30-second epochs, according to the AASM criteria (Version 2.1, 2014) [21, 22]. The resulting, labeled raw data were then used for our standard statistical and superstatistical analysis, and also as a ground truth to test the performance of the Bayesian sleep-stage classification.

### Sleep-stage specific statistical properties of raw EEG data

In a first step, each individual epoch *n* and channel *k* was statistically analyzed by computing the probability density distribution *p*_*n,k*_(*y*) of the momentary EEG signal amplitudes *y*_*n,k*_(*t*), their temporal auto-correlation function

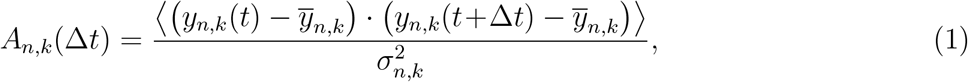

as well as the cross-correlation function between channels 1 and 2

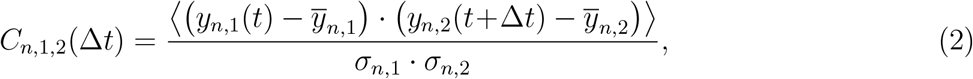

where 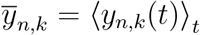 is the temporal average of channel *k*’s amplitude *y*_*n,k*_(*t*) within epoch *n*, and 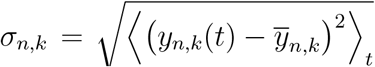 is the corresponding standard deviation. Note that in this case, the standard deviation is equivalent to the root-mean-squared amplitude values that we used in previous studies [8, 9].

In a second step, we have pooled and averaged *p*_*n,k*_(*y*), *A*_*n,k*_(Δ*t*) and *C*_*n*,1,2_(Δ*t*) over all epochs that belong to the same sleep stage *s*. The quantities *p*_*n,k*_(*y*) and *A*_*n,k*_(Δ*t*) were additionally pooled and averaged over all channels *k*. As a result, we obtain the statistical properties *p*_*s*_(*y*), *A*_*s*_(Δ*t*) and *C*_*s*,1,2_(Δ*t*) that are characteristic for each sleep stage *s* and which are shown in Fig. 1.

**Figure 1:**
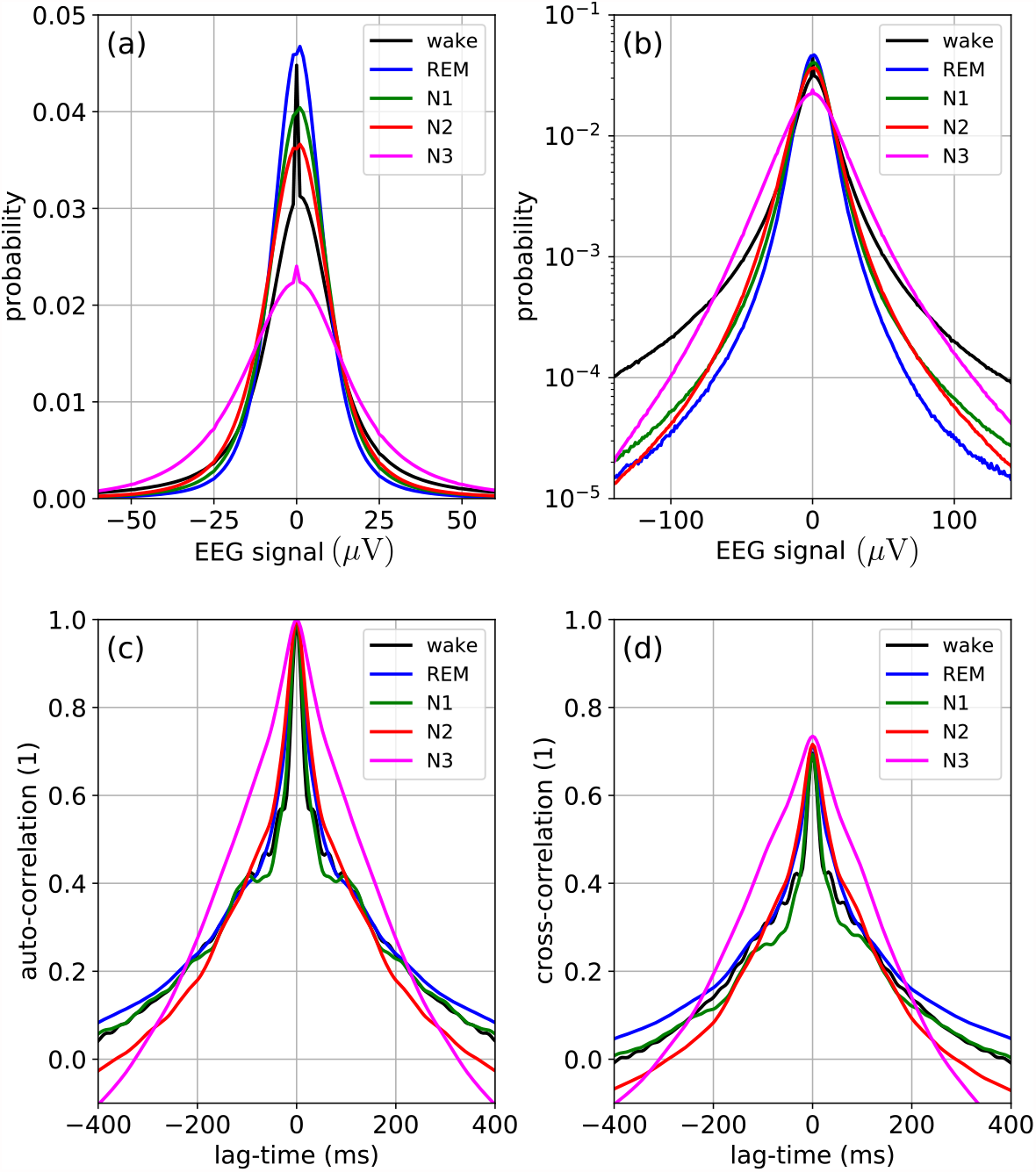
Statistical properties of the raw EEG signals. (a) Linear plot of the probability density distribution. (b) Semi-logarithmic plot of the probability density distribution. (c) Auto-correlation function. (d) Cross-correlation function between channels 1 and 2.

### Extraction and statistical analysis of hyper-parameters

Based on the raw data *y*_*n,k*_(*t*), we have computed for each channel *k* and epoch *n* a set of hyper-parameters, namely the standard deviation

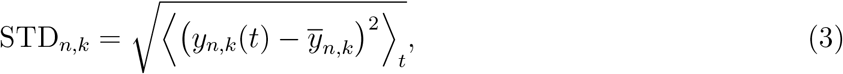

the excess curtosis

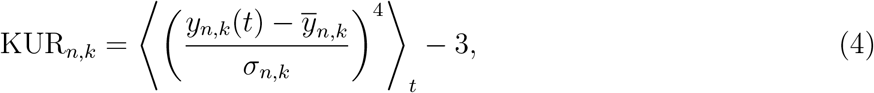

the skewness

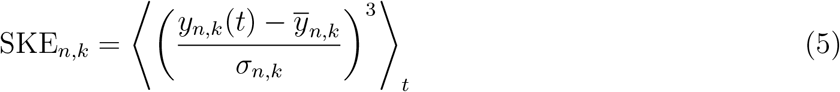

and the value of the auto-correlation function at a specific, fixed lag-time

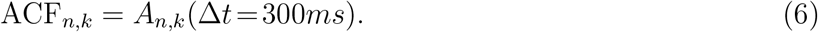

As these hyper-parameters are strongly fluctuating themselves, we have pooled them over all epochs and channels and computed their sleep-stage specific distribution functions *p*_*s*_(*STD*), *p*_*s*_(*KUR*), *p*_*s*_(*SKE*) and *p*_*s*_(*ACF*), which are shown in Fig. 2.

**Figure 2:**
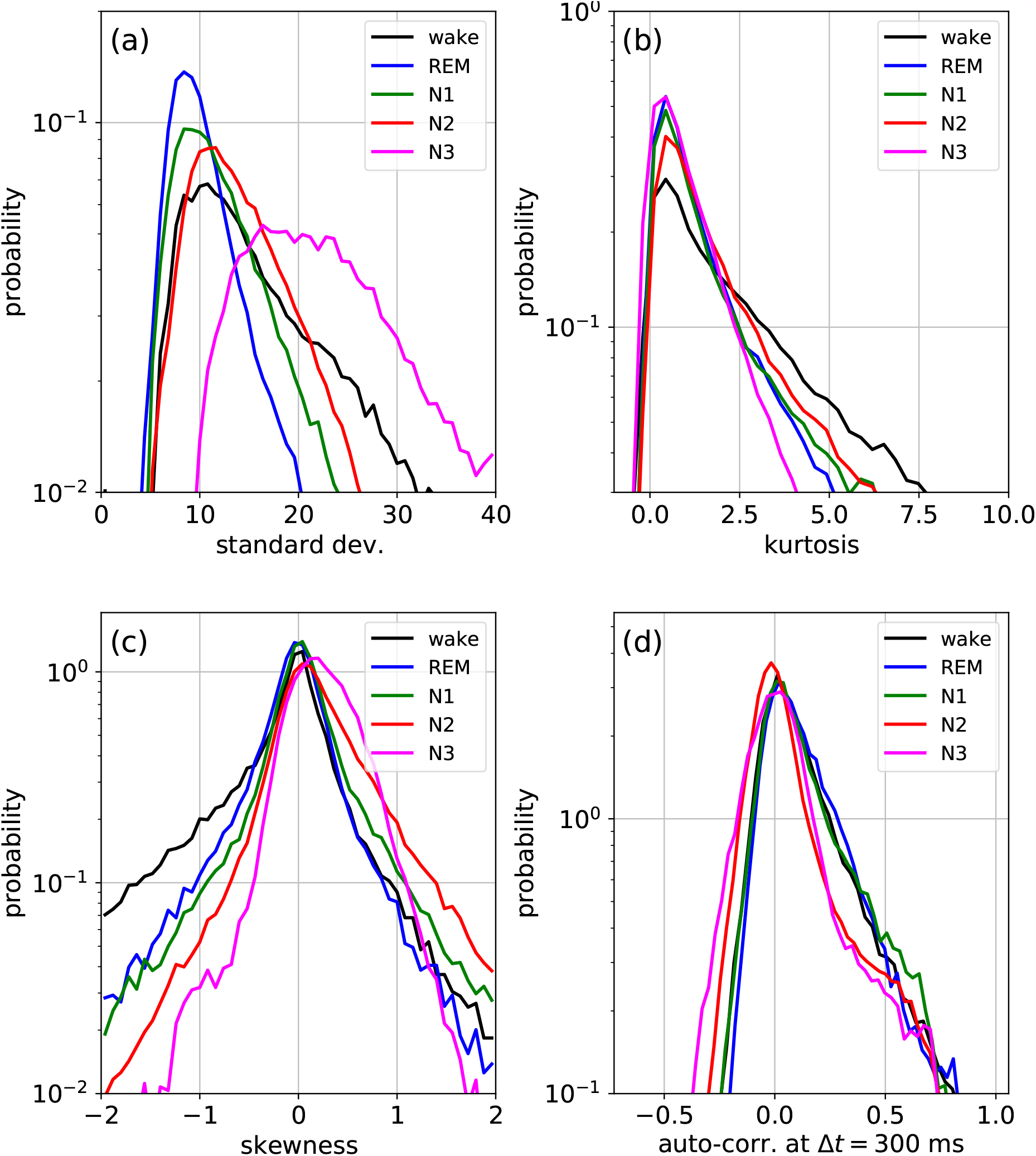
Probability density distributions of hyper-parameters extracted from the raw EEG data shown in Fig.1. (a) Standard deviation STD. (b) Kurtosis KUR. (c) Skewness SKE. (d) Auto-correlation at lag-time 300 ms, denoted as CDT.

### Temporal trend-analysis of hyper-parameters

For a temporal trend-analysis of the hyper-parameters, we no longer partition the EEG time series *y*_*n,k*_(*t*) into 30-second-epochs, but into longer, contiguous ‘sleep phases’: Within a given full-night recording, the sleep phases *J* are defined as the longest possible continuous time periods [*T*_*J,beg*_, *T*_*J,end*_], in which the subject was scored to be in the same constant sleep stage *s* = *s*(*J*). Typically, each sleep phase *J* contains a large number of subsequent epochs *n*. The hyper-parameters STD_*n,k*_, KUR_*n,k*_, … perform a ‘higher-order’ random walk within each sleep phase *J*, and visual inspection reveals that some of these random walks have rising and falling trends (Fig. 3(a)).

**Figure 3:**
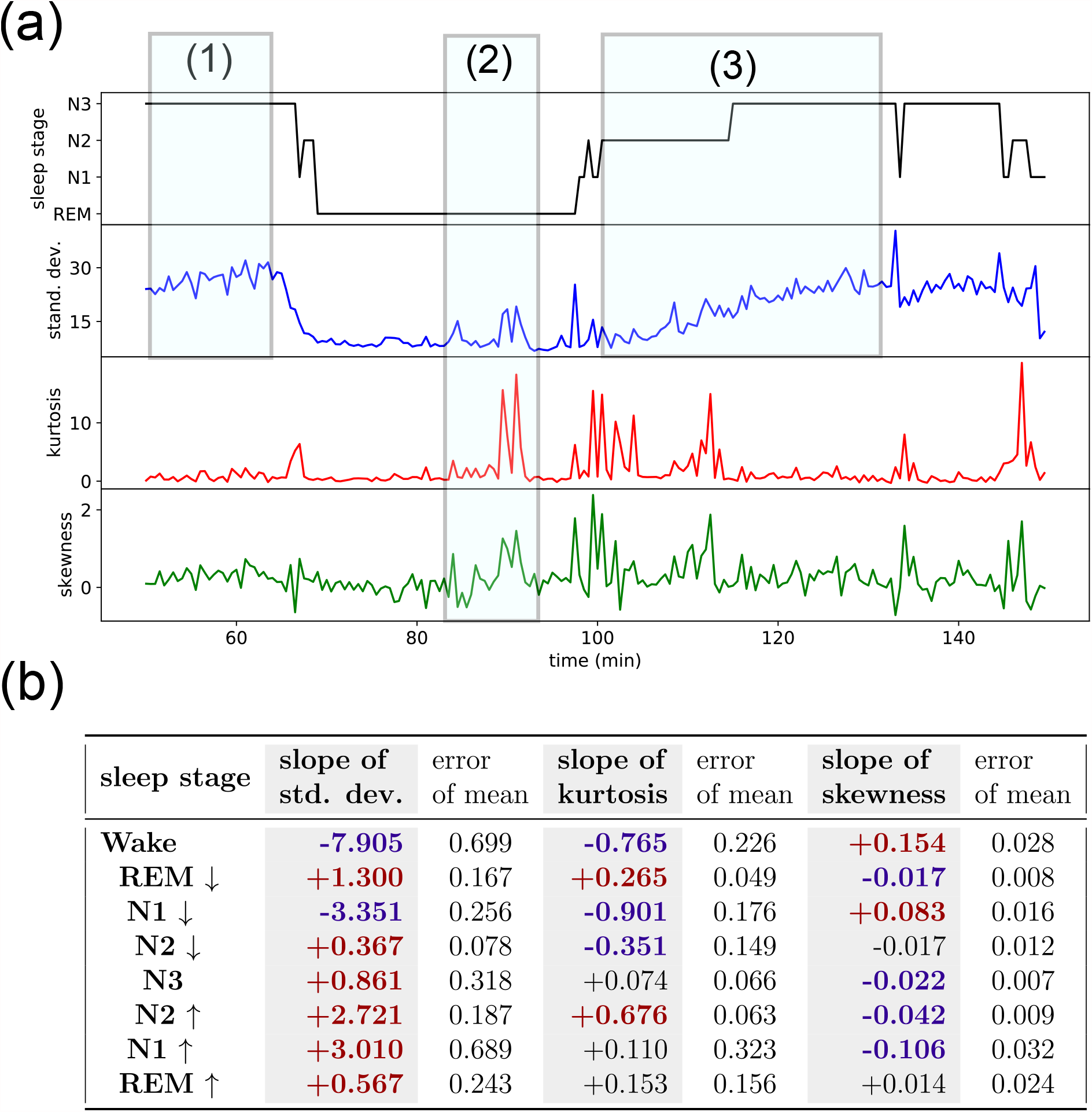
(a) Typical features in the temporal behavior of hyper-parameters, such as consistent trends (1) and extreme fluctuations (2) within an ongoing sleep stage, as well as trends that extend across sleep stages (3). (b) Average slope of hyper-parameters (computed from linear fits) in different sleep stages. The stages REM, N1 and N2 have been separately evaluated for the ‘falling’(↓) and ‘rising’ (↑) phases of the sleep cycle. Some hyper-parameters (bold-face numbers) show a significantly positive or negative sign that is characteristic for certain sleep stages.

To evaluate these trends, we approximate the individual intra-phase time series of the hyper-parameters by linear functions, *f*_*hyp,J*_(*n*)≈*a*_*J*_ ∗ *n* + *b*_*j*_, using least-square fits. The slopes *a*_*J*_ of these linear fits are then pooled and averaged over all sleep phases *J* with the same sleep stage *s*. The results are shown in Fig. 3(b). Note that here we have sub-divided the sleep stages REM, N1 and N2 into the ‘falling’ and the ‘rising’ part of the oscillatory motion between the two ‘extreme’ stages of Wake and N3.

### Evaluation of transition probabilities between sleep stages

The sequence of human-scored sleep stage labels *s*_*n*_ for each subsequent epoch *n* can be regarded as a random walk in a discrete state space *s*_*n*_ ∈ {*Wake, REM, N* 1, *N* 2, *N* 3} . As this discrete random walk shows clear temporal correlations, we have evaluated the (normalized) transition probabilities *p*(*s*_*J*+1_ |*s*_*J*_) between subsequent sleep phases, as well as the transition probabilities *p*(*s*_*n*+1_ |*s*_*n*_) between subsequent epochs. The resulting transition matrices are shown in Fig. 4(a). Note that, by definition, the diagonal elements of the phase-to-phase transition matrix are zero. By contrast, the diagonal elements of the epoch-to-epoch transition matrix are relatively close to one, as each sleep stage has a high degree of persistence.

**Figure 4:**
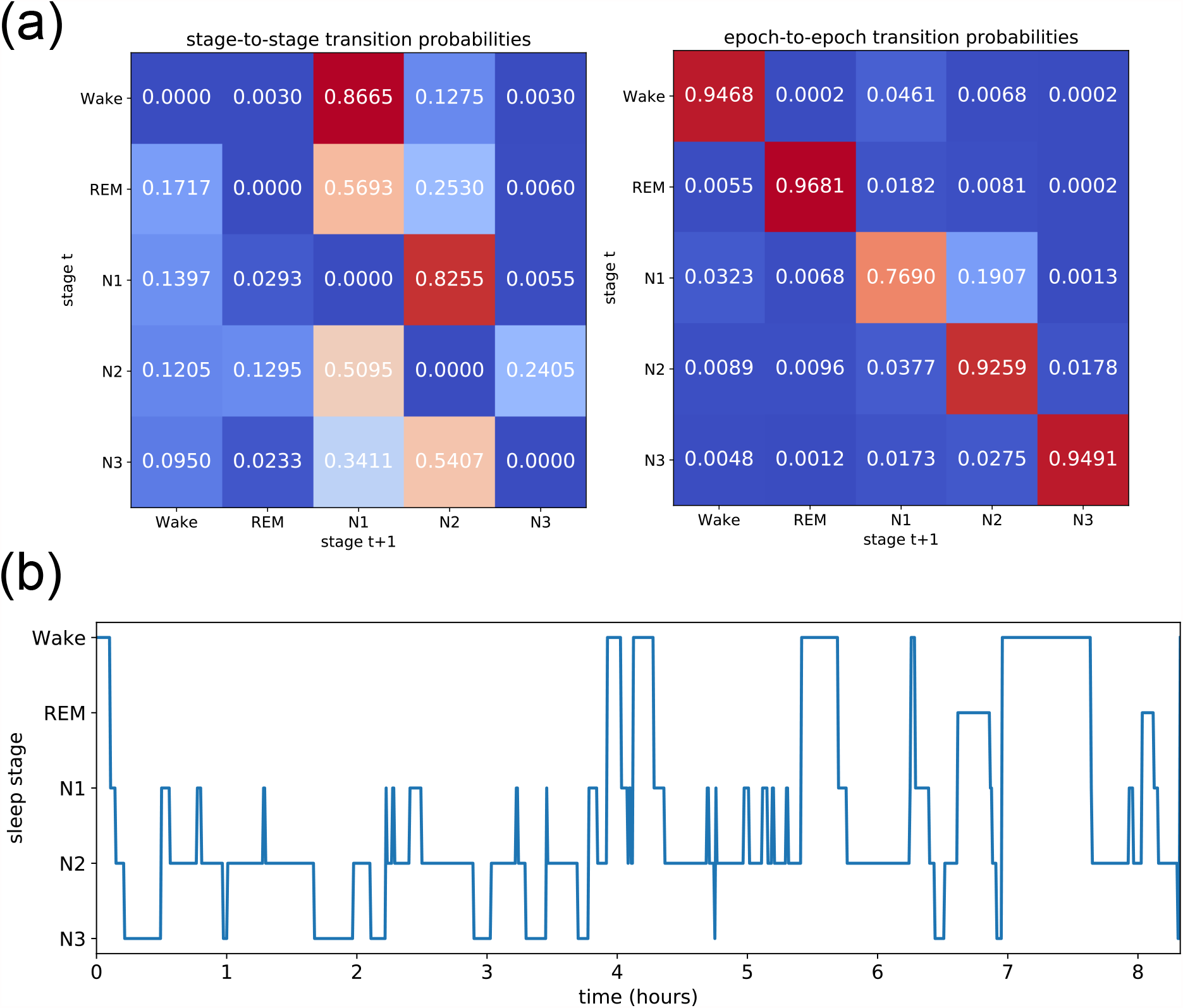
(a) Transition probabilities (color coded) from one non-interrupted sleep stage to the next (top left), and from one 30-second-epoch to the next (top right). The stage-to-stage probabilities describe a strong propensity for transitions from Wake to N1, and from there to N2. After this, the most likely behavior is an oscillation between N1 and N2, or between N2 and N3. (b) Example of a simulated hypnogram, where the random walk between sleep stages is modeled as a Markov process, based on the epoch-to-epoch transition probabilities (top right).

The epoch-to-epoch transition matrix defines a Markov random process of first order. After defining the starting stage *s*_*n*=0_, the transition matrix can be used to simulate an arbitrarily long sequence of sleep stages. An example is shown in the hypnogram of Fig. 4(b).

### Bayesian sleep stage prediction

We have implemented a simple Bayesian model that predicts the probabilities *P* (*s*_*n*_) of the sleep labels *s*_*n*_ ∈ {*Wake, REM, N* 1, *N* 2, *N* 3} from the raw EEG data *D*_*n*_ in each 30-second-epoch *n* (Note that *D*_*n*_ here stands for the complete set of 30*256 successive EEG values corresponding to the given epoch *n*). The prediction is based on the momentary values *h*_*kn*_ of selected statistical hyper-parameters (In our case, the standard deviation *h*_1*n*_, the kurtosis *h*_2*n*_, and the skewness *h*_3*n*_), which are calculated directly from the data *D*_*n*_, and which have different likelihoods *q*(*h*_*kn*_|*s*_*n*_) in the various sleep stages *s*_*n*_. Furthermore, we take into account the prior probability Π(*s*_*n*_) of the momentary sleep stage, which depends on the prediction *P* (*s*_*n*−1_) from the last epoch and on the known transition probability *M* (*s*_*n*_|*s*_*n*−1_). The prediction for the current epoch is then given by

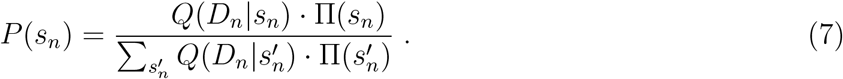

Here, the global likelihood *Q*(*D*_*n*_|*s*_*n*_) of the current data epoch *D*_*n*_ is given as the product over the individual likelihoods of the different hyper-parameters *h*_*kn*_:

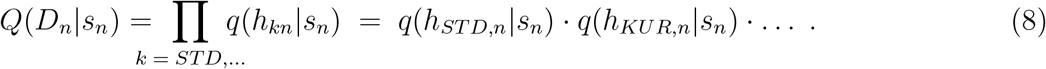

We have numerically implemented these likelihood distribution as continuous spline-extrapolations that were pre-computed from empirical histograms with discrete bins. In this way, also new data can be handled with extreme values of the hyper-parameters that are outside of the empirical histograms. Another possible implementation would be via kernel density distributions. The (normalized) prior probability is computed as

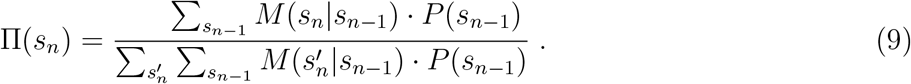

In the initial epoch *n*_0_, we assume for simplicity that the subject is in the wake state. For occasional epochs in which the raw EEG data are not reliable due to obvious measurement artefacts, Bayesian updating proceeds only on the basis of the prior Π.

## 3 Results

Based on the methods described in the section above, we have analyzed the EEG signals on three different time scales: The shortest scale, corresponding to the individual **time steps** *t* in the raw signals (recorded with intervals of approximately 4 milliseconds), the medium scale of **epochs** *n* (each with a duration of 30 seconds), and the longest scale of **sleep phases** *J* (defined as sets of subsequent epochs, in which the subject was scored to be in the same sleep stage *s*). Note that, in contrast to time steps and epochs, sleep phases are time periods with a variable duration.

Our first goal was to establish that single-channel EEG signals *y*_*k*_(*t*) during sleep can be considered as heterogeneous random walks, that is, as random processes in which the statistical properties change over time, and in particular differ between sleep stages. We have therefore analyzed, for each sleep stage *s*, the probability distributions *p*_*s*_(*y*) of the raw EEG amplitudes, its auto-correlation function *A*_*n,k*_(Δ*t*), and the cross-correlation function *C*_*n*,1,2_(Δ*t*) between channels F4-M1 and C4-M1 (Fig. 1). The cross-correlations for the other two combinations of channels did not show qualitatively different results (data not shown).

We find that the amplitude distributions *p*_*s*_(*y*) are non-Gaussian and clearly leptocurtic (positive excess kurtosis) for all sleep stages, an anomaly frequently found in complex systems with superstatistical parameter changes [18]. The distributions for sleep stages REM, N1 and N2 are relatively similar to each other, but the wake and N3 stages are considerably broader. For all sleep stages, the EEG amplitudes show positive temporal auto-correlations up to lag-times of 300 ms, and very similar results are found for the cross-correlation functions. Interestingly, the N3 stage again differs from the other sleep stages, in that it has significantly stronger correlations for lag-times shorter than about 200 ms.

Having thus established the heterogeneous character of sleep EEG recordings, we turn to a super-statistical analysis and study the properties of selected hyper-parameters derived from the raw signals *y*_*k*_(*t*). In particular, we consider as hyper-parameters the standard deviation of the amplitude distribution *p*_*s*_(*y*), its excess kurtosis, its skewness, as well as the value of the auto-correlation function at a specific lag-time Δ*t* = 300ms. We compute the probability distributions of these hyper-parameters in different sleep stages (Fig. 2) and find that they cannot be regarded as approximately constant, as would be the case in a stationary, temporally homogeneous random walk. Instead, the hyper-parameter distributions differ strongly between sleep stages, and their wide distribution within each given sleep stage points to ongoing dynamical changes of brain activity that happen not only at the transition points to new sleep stages, but continuously.

By inspecting the temporal evolution of the hyper-parameters (Fig. 3(a)), we repeatedly observe extreme ‘bursts’ that exceed the normal range of fluctuations. Moreover, we frequently find that certain hyper-parameters rise or fall consistently within and also across sleep phases. A quantitative trend analysis including all available data indeed confirms these anecdotal observations and shows that, on average, some hyper-parameters have statistically significant positive or negative trends that are characteristic for each sleep stage (Fig. 3(b)).

It is also possible to consider sleep as a random walk through the discrete state space of the five sleep stages (Note that, for simplicity, our use of the term ‘sleep stages’ is also including the wake state throughout this work). In this context, we have computed the transition probabilities between subsequent sleep phases and between subsequent epochs, which are presented in the form of 5×5 transition matrices in Fig. 4(a).

The resulting elements of the phase-to-phase transition matrix (left) show that every stage has a preferred successor stage (that is, each row in the matrix has a clear maximum entry). This leads to the emergence of a ‘default’ sequence of stages: Wake →N1→N2. After this, an ongoing oscillation N2↔N1 is most probable, followed by a final ‘decent’ to N3, and eventually the subject will again ‘rise up’ towards the next wake state. In the case of the epoch-to-epoch transition matrix (right), all diagonal elements are close to one, reflecting the strong temporal persistence of each sleep stage.

The epoch-to-epoch transition matrix can be directly used to construct a first-order Markov model for the stochastic succession of sleep stages. Using such a model, an arbitrary number of simulated ‘hypnograms’ can be sampled, and a typical example is shown in Fig. 4(b). Moreover, it may be possible to define a quantitative measure of sleep quality, based on the 25 entries of an individual’s epoch-to-epoch transition matrix, compared to the corresponding values in a reference group of healthy sleepers.

The fact that each sleep stage has a specific distribution of hyper-parameters (compare again Fig. 2) does not only confirm the heterogeneous, non-stationary character of the full-night EEG signals, but it can also be exploited for an automated sleep stage detection. As a proof-of-concept, we have implemented a simple Bayesian sleep stage detector which uses the epoch-to-epoch transition matrix as a prior and the hyper-parameter distributions as likelihood factors. Although the selection of hyper-parameters is arbitrary and the detector has not been optimized in any way, the predictions of the detector are in some cases very close to the ground truth of the human somnologist (Fig. 5(a,b)). Moreover, the distribution of prediction accuracies (defined as the fraction of correctly classified epochs) systematically shifts to larger values, i.e. prediction performance becomes better, when more hyper-parameters are included into the Bayesian likelihood.

**Figure 5:**
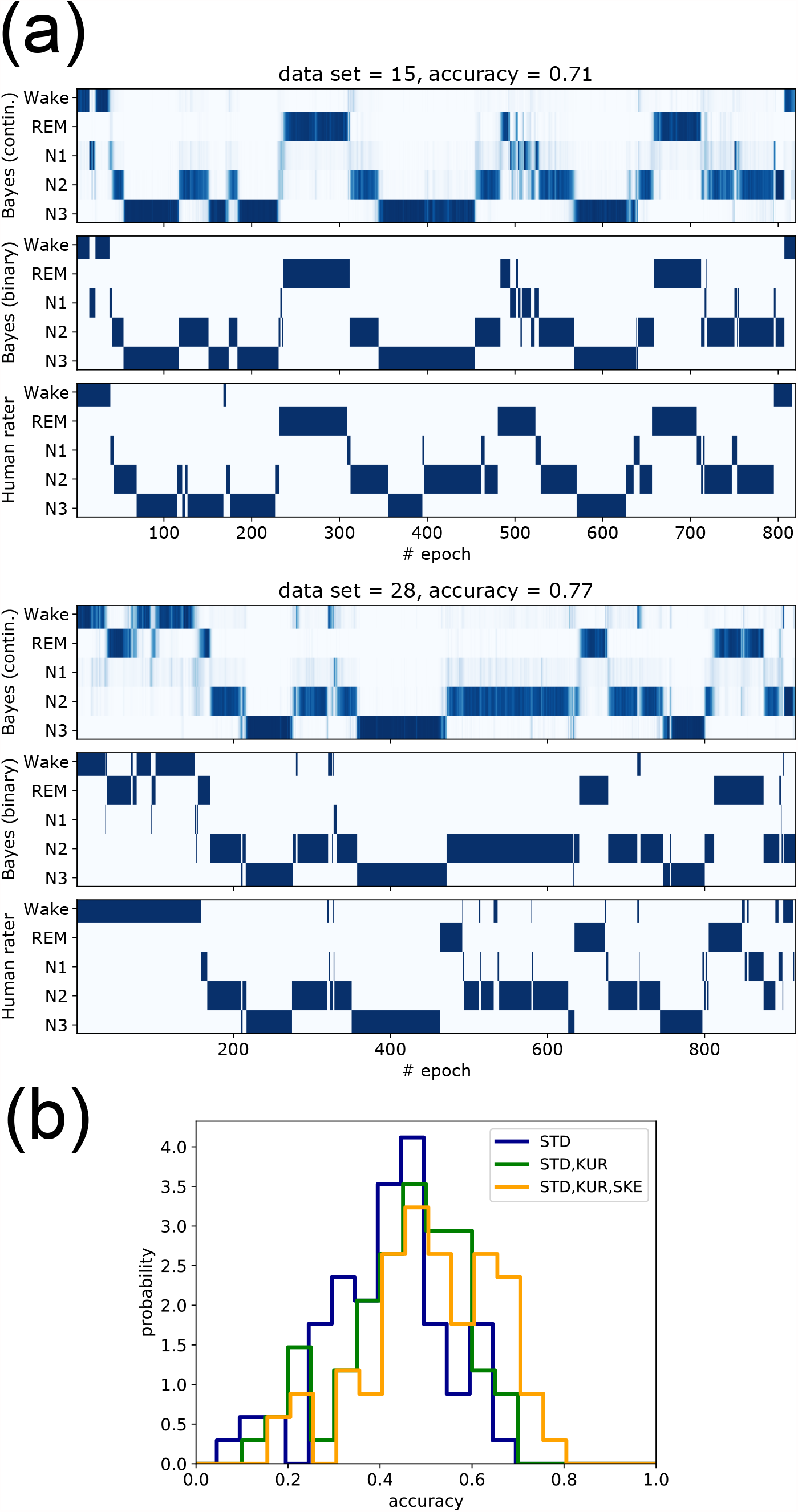
(a) Two examples of automated Bayesian sleep stage classification. In each case, the upper hypnogram shows, for each 30-second epoch, the posterior probabilities of the sleep stages, with larger color density corresponding to larger probability. The middle hypnogram shows only the predicted sleep stage with maximum posterior probability. The lower hypnogram is the ground truth, provided by the specialist human rater. The accuracies are defined as the ratio of correct sleep stage predictions. (b) Distributions of accuracies over all 68 data sets, with different combinations of hyper-parameters used in the Bayesian likelihood. The average prediction quality can be improved by simply adding more hyper-parameters.

## 4 Discussion

Traditionally, the analysis of EEG recordings has mainly focused on the oscillatory features of the signals, such as alpha-, beta-, delta- and theta-frequency bands, and in the context of sleep also on wavelet-like features, i.e. grapho-elements, such as sleep spindles or K-complexes. Just recently it became clear that also the aperiodic component of an EEG signal, in particular the ubiquitous ‘background noise’ with a *f* ^−*β*^-like power-spectrum, contains valuable information about the physiological state of the subject [23, 24]. Indeed, these aperiodic, scale-free fluctuations have been shown to systematically change with age and with the tasks to be performed [25]. Moreover they offer an alternative way to asses the level of arousal [26].

In this work, we have investigated an alternative approach that is neither based on oscillatory features, nor on the global power spectrum of the EEG signal. Instead, we treat the signal as a non-stationary, heterogeneous random walk, generated by a stochastic system with parameters that change over time, depending on the physiological state of the subject.

In particular, this random walk has different statistical properties in each of the five sleep-(or, more precisely, vigilance-) stages, and these differences can be exploited for a simple automated Bayesian sleep stage detection. As a proof-of-concept, we have implemented a first version of such a detector and demonstrated that it achieves relatively high detection accuracies. In contrast to sleep stage detectors based on deep neural networks which suffer from the ‘black box problem’ [27], our Bayesian approach is completely transparent and explainable, as the features used to distinguish between sleep stages (i.e. the distributions of hyper-parameters) are explicit. Once these hyper-parameter distributions are extracted from the raw data and included into the likelihood, the Bayesian detector can immediately be applied without any training or further optimization. In contrast, most deep learning applications require extensive training and are ‘data hungry’ [28]. Furthermore, while the posterior probabilities of the momentary sleep stages are mathematically well-defined in the Bayesian approach, it is not clear if the typical softmax outputs of a deep neural network can actually be interpreted as probabilities, or if they are just a list of scores that sum up to one. Finally, we have shown that the accuracy of the Bayesian sleep stage detector can be systematically improved, simply by including additional hyper-parameter distributions as factors in the likelihood. In principle, the number of these factors could be made arbitrarily large by using hyper-parameters such 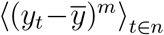the *m*-th central moments of the fluctuating EEG signal *y*_*t*_ within each 30-second epoch *n*, or 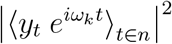the magnitude squared of momentary Fourier components for different frequencies *ω*_*k*_.

Besides the probability distributions of the hyper-parameters, we have also studied their gradual evolution over time. Some of the hyper-parameters show consistent rising or falling trends within and across the phases in which the subject is scored to be in a constant sleep stage. For example, we have observed a case where the standard deviation of the EEG signal is continuously increasing for about 30 minutes, while the subject is passing from the REM state through N1, to N2, and finally to the N3 state (compare Fig. 3(a)(3)). Such a gradual buildup of EEG amplitude points to a continuous mechanism in the brain that is regulating the sleep cycle, a phenomenon similar to the change of hyper-parameters that we have observed in migrating cells during the cell cycle [18].

We speculate that, rather than sub-dividing sleep into discrete stages, it might be useful to introduce a continuous ‘master-variable’ *ϕ*(*t*) ∈[0, 2*π*], roughly resembling the mathematical phase of a sinusoidal oscillation, which tends to increase about linearly with time and which reflects the momentary position of the subject within the sleep cycle. In principle, it may then be possible to design a simple stochastic ‘first-level’ model, such as an auto-regressive process of low order, with coefficients that are not constant but which are top-down controlled (from a second model level) by the master variable *ϕ*(*t*). As we have demonstrated in other contexts [18, 19], such super-statistical two-level models are often capable to reproduce the anomalous time-dependent statistics of biological and other complex systems (typically involving non-normally distributed, long-time-correlated signals) in a particular simple way. In future work, one could therefore attempt to reproduce the sleep-stage dependent properties of the EEG raw signals (Fig. 1) and of the various hyper-parameters (Fig. 2) with such a two-level model.

Indeed, it may even be possible to relate the phase variable *ϕ*(*t*) to existing models of sleep [29]. An obvious candidate would be the famous two-process model [30, 31], in which sleep is controlled by the non-linear interplay between the circadian propensity for sleep, governed by an intrinsic circadian oscillator, and a homeostatic drive for sleep that continuously increases during the waking state and dissipates during sleep. In this case, the circadian and homeostatic signals may directly represent the second-level control signals of a two-level super-statistical model.

## 5 Additional information

### Author contributions statement

CM has conceived of applying super-statistical analysis to sleep data, implemented the methods, evaluated the data, and wrote the paper, PK designed the study, discussed the results and wrote the paper, AS discussed the results, MT provided access to resources and wrote the paper, HS provided access to resources and wrote the paper.

## Data availability statement

Data and analysis programs will be made available upon reasonable request.

## Ethical approval and informed consent

The study was conducted in the Department of Otorhinolaryngology, Head Neck Surgery, of the Friedrich-Alexander University Erlangen-Nürnberg (FAU), following approval by the local Ethics Committee (323 – 16 Bc). Written informed consent was obtained from the participants before the cardiorespiratory poly-somnography (PSG).

## Third party rights

All material used in the paper are the intellectual property of the authors.

## References

[1] Witold Pedrycz and S Chen. Time series analysis, modeling and applications. A Computational Intelligence Perspective (e-book Google), 2013.

[2] Gilles Zumbach. Discrete time series, processes, and applications in finance. Springer Science & Business Media, 2012.

[3] Gebhard Kirchgässner, Jürgen Wolters, and Uwe Hassler. Introduction to modern time series analysis. Springer Science & Business Media, 2012.

[4] Ali Bulent Cambel. Applied chaos theory: A paradigm for complexity. Elsevier, 1993.

[5] Theodore L Baker. Introduction to sleep and sleep disorders. Medical Clinics of North America, 69(6):1123–1152, 1985.

[6] James D Geyer, Sachin Talathi, and Paul R Carney. Introduction to sleep and polysomnography. Clinical Sleep Disorders. Philadelphia: Lippincott Williams & Wilkins, pages 265–266, 2009.

[7] John S Barlow. The electroencephalogram: its patterns and origins. MIT press, 1993.

[8] Patrick Krauss, Achim Schilling, Judith Bauer, Konstantin Tziridis, Claus Metzner, Holger Schulze, and Maximilian Traxdorf. Analysis of multichannel eeg patterns during human sleep: a novel approach. Frontiers in human neuroscience, 12:121, 2018.

[9] Patrick Krauss, Claus Metzner, Achim Schilling, Konstantin Tziridis, Maximilian Traxdorf, Andreas Wollbrink, Stefan Rampp, Christo Pantev, and Holger Schulze. A statistical method for analyzing and comparing spatiotemporal cortical activation patterns. Scientific reports, 8(1):1–9, 2018.

[10] Maximilian Traxdorf, Patrick Krauss, Achim Schilling, Holger Schulze, and Konstantin Tziridis. Microstructure of cortical activity during sleep reflects respiratory events and state of daytime vigilance. Somnologie, 23(2):72–79, 2019.

[11] Achim Schilling, Andreas Maier, Richard Gerum, Claus Metzner, and Patrick Krauss. Quantifying the separability of data classes in neural networks. Neural Networks, 2021.

[12] Bradley V Vaughn and Peterson Giallanza. Technical review of polysomnography. Chest, 134(6):1310–1319, 2008.

[13] Joseph W Burns, Leslie J Crofford, and Ronald D Chervin. Sleep stage dynamics in fibromyalgia patients and controls. Sleep Medicine, 9(6):689–696, 2008.

[14] Patrick Krauss, Claus Metzner, Nidhi Joshi, Holger Schulze, Maximilian Traxdorf, Andreas Maier, and Achim Schilling. Analysis and visualization of sleep stages based on deep neural networks. Neurobiology of sleep and circadian rhythms, 10:100064, 2021.

[15] Christian Beck and Ezechiel GD Cohen. Superstatistics. Physica A: Statistical mechanics and its applications, 322:267–275, 2003.

[16] EGD Cohen. Superstatistics. Physica D: Nonlinear Phenomena, 193(1-4):35–52, 2004.

[17] Christian Beck. Generalized statistical mechanics for superstatistical systems. Philosophical Transactions of the Royal Society A: Mathematical, Physical and Engineering Sciences, 369(1935):453–465, 2011.

[18] Claus Metzner, Christoph Mark, Julian Steinwachs, Lena Lautscham, Franz Stadler, and Ben Fabry. Superstatistical analysis and modelling of heterogeneous random walks. Nature communications, 6(1):1–8, 2015.

[19] Christoph Mark, Claus Metzner, Lena Lautscham, Pamela L Strissel, Reiner Strick, and Ben Fabry. Bayesian model selection for complex dynamic systems. Nature communications, 9(1):1– 12, 2018.

[20] William O Tatum, Barbara A Dworetzky, and Donald L Schomer. Artifact and recording concepts in eeg. Journal of clinical neurophysiology, 28(3):252–263, 2011.

[21] Conrad Iber. The aasm manual for the scoring of sleep and associated events: Rules. Terminology and Technical Specification, 2007.

[22] American Academy of Sleep Medicine et al. The aasm manual for the scoring of sleep and associated events: rules, terminology and technical specifications, version 2.0. American Academy of Sleep Medicine, 2012.

[23] Biyu J He, John M Zempel, Abraham Z Snyder, and Marcus E Raichle. The temporal structures and functional significance of scale-free brain activity. Neuron, 66(3):353–369, 2010.

[24] Biyu J He. Scale-free brain activity: past, present, and future. Trends in cognitive sciences, 18(9):480–487, 2014.

[25] Thomas Donoghue, Matar Haller, Erik J Peterson, Paroma Varma, Priyadarshini Sebastian, Richard Gao, Torben Noto, Antonio H Lara, Joni D Wallis, Robert T Knight, et al. Parameterizing neural power spectra into periodic and aperiodic components. Nature neuroscience, 23(12):1655–1665, 2020.

[26] Janna D Lendner, Randolph F Helfrich, Bryce A Mander, Luis Romundstad, Jack J Lin, Matthew P Walker, Pal G Larsson, and Robert T Knight. An electrophysiological marker of arousal level in humans. Elife, 9:e55092, 2020.

[27] Davide Castelvecchi. Can we open the black box of ai? Nature News, 538(7623):20, 2016.

[28] Gary Marcus. Deep learning: A critical appraisal. arXiv preprint 1801.00631, 2018.

[29] John H Abel, Kimaya Lecamwasam, Melissa A St Hilaire, and Elizabeth B Klerman. Recent advances in modeling sleep: from the clinic to society and disease. Current Opinion in Physiology, 15:37–46, 2020.

[30] Alexander A Borbély. A two process model of sleep regulation. Hum neurobiol, 1(3):195–204, 1982.

[31] Serge Daan, DG Beersma, and Alexander A Borbély. Timing of human sleep: recovery process gated by a circadian pacemaker. American Journal of Physiology-Regulatory, Integrative and Comparative Physiology, 246(2):R161–R183, 1984.

